# Probabilistic Brain Extraction in MR Images via Conditional Generative Adversarial Networks

**DOI:** 10.1101/2022.03.14.484346

**Authors:** Saeed Moazami, Deep Ray, Daniel Pelletier, Assad A. Oberai

## Abstract

Brain extraction, which refers to the task of segmenting brain tissue in an MR image of a subject, forms an essential first step for many quantitative neuroimaging applications. These include quantifying grey and white matter volumes, monitoring neurological diseases like multiple sclerosis (MS) and Alzheimer’s disease, and estimating brain atrophy. Over the years several algorithms that automate brain extraction have been proposed. More recently, novel image-to-image deep learning methods have been implemented for this task, and have demonstrated significant gains in accuracy and robustness. However, to our knowledge, none of these algorithms account for the uncertainty that is inherent in brain extraction. Motivated by this, we propose a novel, probabilistic deep learning algorithm for brain extraction that recasts this task as a Bayesian inference problem, and then utilizes a conditional generative adversarial network (cGAN) to solve it. The input to the generator network is an MR image of the head and the output is a collection of images of the brain that are drawn from a probability density conditioned on the input image. These images are used to generate a pixel-wise mean image, which serves as the best guess for an image of the brain, and a pixel-wise standard deviation image, which quantifies the uncertainty in the prediction. We test this algorithm on head MR images of fifty subjects and demonstrate that it is more accurate than a commonly used brain extraction tool, and that its performance compares well with the current state of the art in deep learning algorithms. We also demonstrate the utility of the estimates of uncertainty generated by the algorithm.

## 1 Introduction

MRI brain extraction, or skull stripping, is the process of segmenting brain parts, namely the cerebrum, cerebellum, and brain stem organs, in a whole head MR image. The output of this task is usually in the form of a 3D image volume of the brain with the complimentary parts eliminated, or a 3D binary mask volume, which distinguishes a unified brain from the background voxels. The resulting extracted brain images can be used directly by the end-user, or by downstream tools in medical imaging applications, which include grey and white matter volume measurement, monitoring of neurological diseases such as multiple sclerosis (MS) and Alzheimer’s disease, brain atrophy estimation, and brain lesion segmentation. The breadth of these applications makes brain extraction one of the most commonly performed tasks in quantitative brain neuroimaging. Consequently, the accuracy and the confidence with which this task is performed significantly impact subsequent tasks.

Given that skull stripping forms the basis of a wide range of applications, over the years many tools have been developed to automate and optimize this task (see Section 2). However, there are still several aspects that can be improved. These include further gains in accuracy, robustness (the ability to work with heterogeneous and diverse MR images), and speed, especially for high throughput applications. They also include the ability to quantify the uncertainty in skull stripping. That is, providing the end-user with the best estimate of the brain along with measures of confidence in that estimate. The method proposed in this manuscript addresses and makes gains in all these challenging aspects of skull stripping.

This paper introduces a novel deep Bayesian inference framework in which the brain extraction task is performed using a conditional generative adversarial network (cGAN) [1] architecture. The cGAN model is trained using a set of pairwise MR images of the head (denoted by ***h***) and the corresponding extracted brain (denoted by ***b***). Then using Bayes’ theorem, the model learns the distribution of the brain image conditioned on the corresponding image of the head, that is *p*_*B*|*H*_ (***b***|***h***). Thereafter, given any new MR image of the head, the model is able to efficiently sample from the conditional distribution. These samples, in turn, can be used to calculate important statistics of the distribution, including the pixel-wise mean and standard deviation. The former provides an estimate of the extracted brain image, while the latter quantifies the uncertainty in the extraction, indicating the model’s confidence in generating different regions of the extracted brain. This information can be used by the end-user in downstream applications. For example, they may focus on regions with higher uncertainty for potential quality control or use the cumulative measure of standard deviation as a surrogate of the estimated error in the extractions for the entire brain. Our experiments show that the proposed algorithm outperforms the state-of-the-art methods for MRI brain extraction in terms of accuracy and robustness, and does so in acceptable computational time. Additionally, to the best of our knowledge, it is the only method that provides an estimate of uncertainty in its prediction.

The rest of the paper is organized as follows. In Section 2, a brief review of the related work in this area is provided. In Section 3, the proposed method is described in detail. Numerical results are presented in Section 4, followed by the conclusions and directions for future research in Section 5.

## 2 Related Work

Manual segmentation of the brain in MR images is a labor-intensive and time-consuming task. This has led to a variety of automated methods devoted to solving this problem. In review papers (see [2, 3], for example), brain extraction tools are categorized as conventional, machine learning (ML)-based, and deep learning (DL)-based methods, with subcategories within each class. Despite the growth of interest in ML/DL methods, some tools based on conventional methods are still widely used, especially in medical research and clinical environments. These include Fsl [4], BET [5], BEaST [6], and AFNI 3dSkullStrip [7].

More recently, DL-based methods have received attention. They are typically more robust when working with images with intensity variation and slight image misalignment, and therefore do not require extensive pre-processing and parameter adjustment. This robustness extends to working with images other than *T* 1 weighted images that are usually used for brain extraction. Moreover, since they rely on using GPUs, their speed can be readily scaled. Among these methods, those that utilize the U-net architecture [8] have been successful in performing different image-to-image tasks in medical applications [9]. Accordingly, most DL-based brain extraction approaches utilize the U-net architecture. In the following paragraph, we provide a summary of some of these methods.

In the auto-context convolutional neural network (Auto-Net) [10], multiple fully connected and convolutional neural network layers are combined within a U-net structure. Models are then trained to perform brain extraction using images sliced along axial, sagittal, and coronal directions. Multiple 2D patch sizes are used to capture the context of different spatial scales. Segmentation metrics (see Section 3.3) are then evaluated for different subjects. The average dice similarity coefficients for healthy subjects, fetal subjects, and subjects diagnosed with Alzheimer’s disease are reported to be 97.73, 93.80, and 97.62, respectively. In a separate work [11], the performance of a U-net based artificial neural network is compared with a number of conventional tools using different types of MR images. The best results (average dice similarity coefficient of 97.6) are obtained with T1 weighted images. The complementary segmentation network (CompNet) [12] method utilizes two pathways to learn features from both the brain and complementary tissues, i.e., bones and other parts of the head. The two trained models then work together to enhance the extraction accuracy. The model achieves a dice similarity coefficient of 98.27 and sensitivity of 98.26 for healthy subjects and 97.62 and 97.84 for subjects with pathological conditions. Finally, in [13] and [14], the authors investigate 3D U-net structures for brain extraction and report dice coefficients of 99.50 and 95.77, respectively.

The algorithm developed in this manuscript also uses a 2D U-net architecture and attains. However, in contrast to the methods described above, it does so in a probabilistic context. Rather than producing a single image of the brain, it produces an ensemble of images which can then be used to extract the most likely image and to assess the uncertainty in this prediction. The accuracy of the proposed method, as measured by the dice coefficient and other metrics, compares with the very best results reported in the literature.

## 3 Materials and Methods

This section is dedicated to describe the brain extraction task and its formulation as a Bayesian inference problem. The proposed methods, the dataset used to train and test the model, and the evaluation metrics are also explained.

### 3.1 Problem formulation

The brain extraction task is defined as a Bayesian inference problem that is solved using a deep generative adversarial network (GAN) [15]. In particular, given paired sample images of the head and the extracted brain, a conditional GAN (cGAN) is trained to produce samples from the conditional distribution.

The image size of a 3D MR image is denoted by *N*_1_ × *N*_2_ × *N*_3_, where *N*_1_, *N*_2_, and *N*_3_ are the number of voxels in the coronal, sagittal, and axial directions, respectively. An axial slice of the head, denoted by 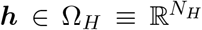, is a two dimensional array of size *N*_*H*_ = *N*_1_ × *N*_2_ pixels. The corresponding image of the brain is denoted by 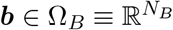, and is of the same size as ***h***, *N*_*B*_ = *N*_*H*_. In the brain image the intensity of all pixels that are not a part of the brain, is set to zero.

It is assumed that a dataset containing *N* pairwise MR images, *S* = {(***h***^(*i*)^, ***b***^(*i*)^)}, *i* = 1, …, *N*, from several subjects is available, and each sample in this dataset is drawn from the joint density function *P*_*BH*_(***b, h***), which itself is unknown. The goal is to utilize this dataset to develop an algorithm for efficiently sampling from the conditional distribution *P*_*B*|*H*_ (***b***|***h***). That is, given a new image of the head, we wish to generate samples images of the brain conditioned on it. Then, we can use these samples to generate the mean brain image, and a standard deviation image, which is used to quantify the uncertainty in the prediction. As described below, this is done via a modified version of the cGAN model [16, 15].

### 3.2 Conditional GAN (cGAN)

As illustrated in Figure 1, the cGAN comprises two deep neural networks, namely, a generator ***g*** and a critic *d*. The generator, ***g*** : Ω_*H*_ × Ω_*Z*_ ↦ Ω_*B*_, accepts as input an image of the head, ***h***, and for any instance of the latent vector 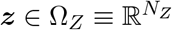 generates an image of the brain. That is, ***b***^*g*^ = ***g***(***h, z***). The latent vector is drawn from the distribution *P*_*Z*_, which is the standard multivariate normal distribution. For a given image of the head, ***h***, by sampling ***z*** from *P*_*Z*_ the generator generates an ensemble of brain images. These can be thought to be drawn from a conditional distribution, 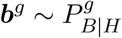. The precise form of this distribution is determined by the weights of the generator, and the goal of the training procedure is to make this distribution as close as possible to the true conditional distribution *P*_*B*|*H*_.

**Figure 1:**
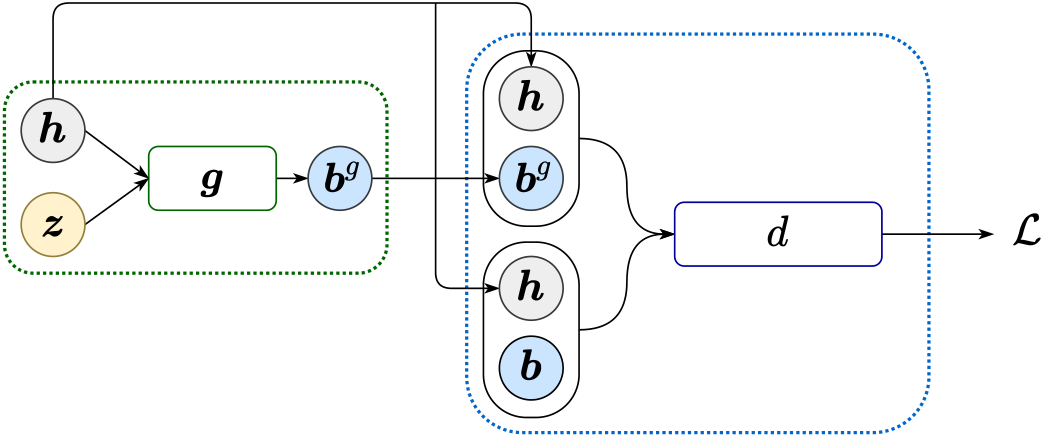
Conditional generative adversarial network (cGAN) architecture.

The critic, defined as *d* : Ω_*H*_ × Ω_*B*_ ↦ ℝ, is responsible for distinguishing between image pairs from the true dataset, i.e., (***h, b***) ∼ *P*_*BH*_, and from the set where the brain image is generated by the generator network, i.e., (***h, b***^***g***^), where ***h*** ∼ *P*_*H*_ and 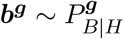. The critic is trained to attain a large value for images from the true dataset and a small value for images generated by the generator. This is done via the Wasserstein GAN loss function [17]:

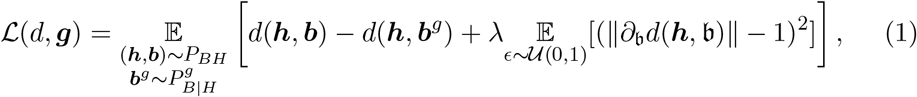

where 𝔼 denotes the expectation. The first two terms on the right hand side of Equation 1 measure the difference between the values of the critic for true and generated images. The last term is the gradient penalty term [18], where *λ* is the gradient penalty coefficient and *𝔟* = *ϵ****b*** + (1 − *ϵ*)***b***^***g***^, where *ϵ* is a random number selected from a uniform distribution 𝒰(0, 1).

The optimal critic and generator, denoted by *d*∗ and ***g***∗, respectively, are given by,

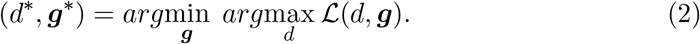

Solving the min-max problem above is equivalent to minimizing the Wasserstein-1 distance between 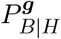 and *P*_*B*|*H*_ [15]. That is, for a trained network, 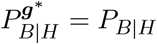 in the Wasserstein-1 distance.

#### 3.2.1 Sampling from the cGAN

Since convergence in the Wasserstein-1 distance implies weak convergence [19], for the trained generator we have,

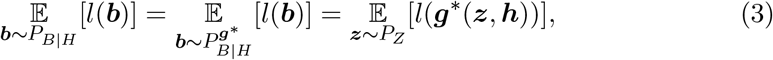

where *l* is any continuous bounded function defined on Ω_*B*_. In other words, for a given image ***h***, computing the expected value of any function of the brain ***b*** over the conditional distribution is the same as computing the expectation over the latent space of the same function applied to images obtained by passing the latent vector through the fully trained generator ***g***∗. Since the dimension of the latent space is typically small (*N*_*Z*_ = 256 in this study), and the cost of forward propagation through the generator network is low, this sampling process is a computationally feasible task to perform.

As depicted in Figure 2, for a given image of the head, the pixel-wise mean image of the brain is calculated using the fully trained generator by sampling the latent vector from its distribution. It is given by,

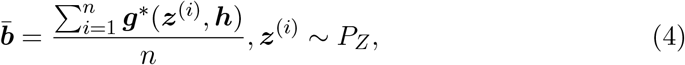

where *n* is the number of samples. Thereafter, an image of the brain volume, denoted by 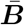 is constructed by stacking these individual slices.

**Figure 2:**
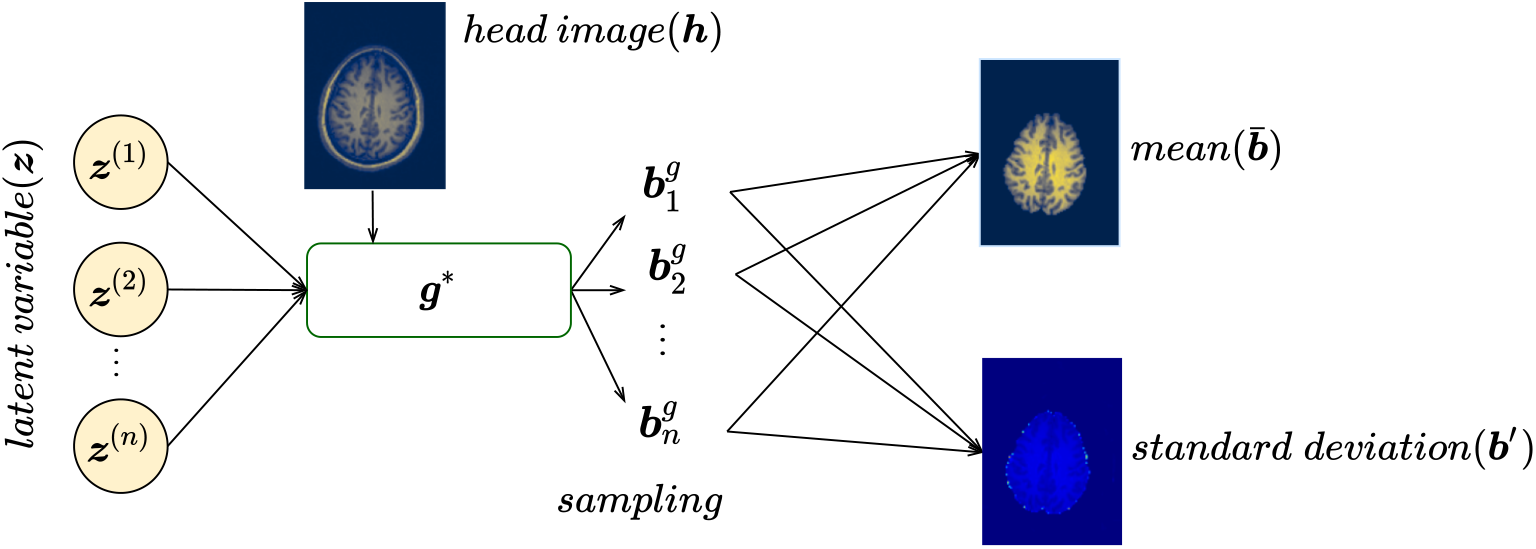
Sampling from the trained generator ***g***∗ to produce mean and standard deviation images.

Similarly, an image of the pixel-wise standard deviation, ***b***′, is calculated by,

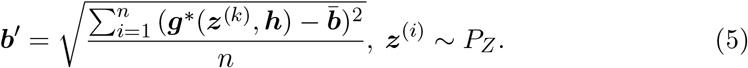

In the equation above the power of 2 is interpreted as a pixel-wise square of an image. The standard deviation image slices are also stacked to yield a volumetric image of pixel-wise standard deviation, denoted by ***B***′.

#### 3.2.2 Post-processing

The final step in this process involves applying a filter to 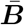 wherein all pixels with intensity greater than a threshold hyper-parameter are set to one and the others are set to zero. The result is a three-dimensional binary image, or the mask, and is denoted by ***M***. In the mask the brain voxels are denoted by 1 and the background is denoted by 0. In the present work, we set the threshold value to 0.04, which is achieved by trial and error.

In the mask generated by the cGAN, we observe some small scattered collection of incorrectly labeled voxels. These occur outside the brain, typically in the lower and upper slices, and near the intersection of the cerebrum and the cerebellum.

To address this issue, we apply two morphological filters to ***M*** that remove islands and cavities smaller than a specified minimal volume (see `remove_small_objects` and `remove_small_holes` operations in [20]). The mask obtained after these filters is not very sensitive to the value of minimal volume parameter as the size of the islands and cavities are much smaller than the size of the brain. Further, in the results section for every subject, we report metrics for mask before and after applying these filters. We note that the mask is very accurate even prior to the application of these filters, and their overall effect is to slightly improve the performance.

#### 3.2.3 Generator and critic architecture

The generator ***g*** is implemented using a deep U-net architecture (see Figure 3). Its input consists of the image of the head, ***h***, and the latent vector, ***z***. It includes a convolution layer, followed by three down-sampling blocks, one central customized residual network (ResNet) block, three up-sampling blocks, and two convolution layers (see Appendix A for a detailed description). As the input is transmitted through the down-sampling blocks, its spatial resolution reduces, while the number of features increases. Exactly the opposite happens in the up-sampling block. Further, information from a given level of spatial resolution is directly transmitted from the down-sampling branch to the up-sampling branch via skip connections. Stochasticity is introduced in the network by utilizing the latent vector ***z*** to perform conditional instance normalization [21] operations at multiple spatial scales. A Sigmoid output function (applied pixel-wise) ensures the predicted brain image is bounded between 0 and 1.

**Figure 3:**
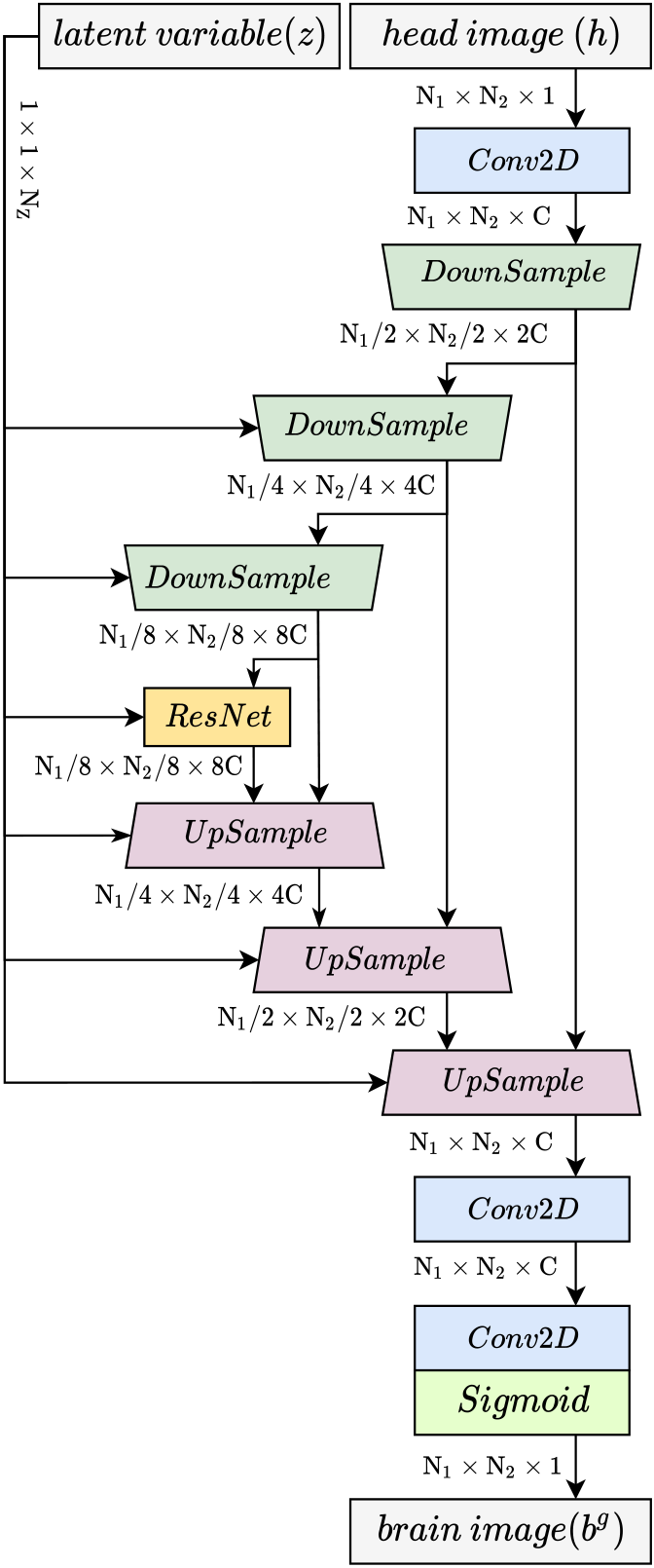
The U-net based architecture of the generator.

The architecture of the critic network is shown in Figure 4. The input is a set of paired images of the brain and the head. The input is passed through a convolution layer followed by three down-sampling and ResNet blocks. Similar to the generator network, down-sampling blocks reduce the spatial size of their input by a factor of two and doubles the number of channels. Further, conditional instance normalization is replaced by layer normalization. The down-sampling blocks are followed by two dense layers, with the final output being a scalar.

**Figure 4:**
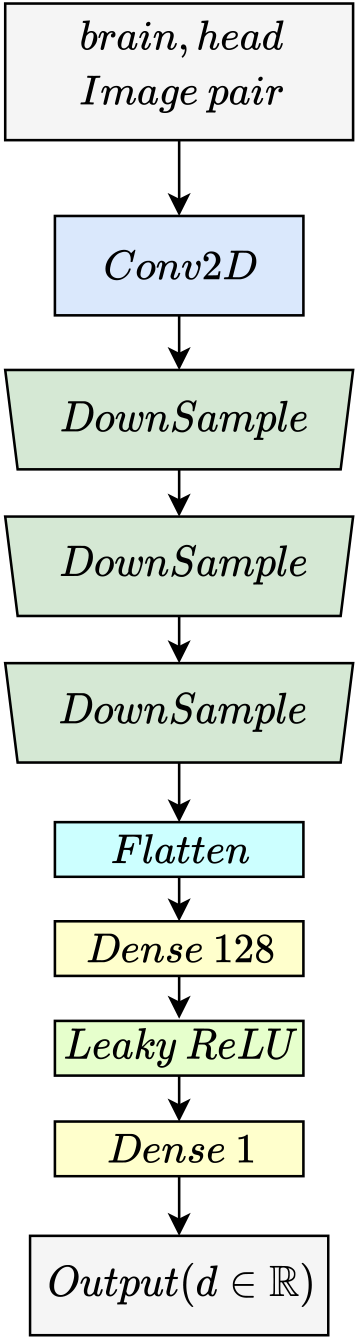
The critic neural network architecture.

The detailed architecture of the generator and critic blocks are presented in Appendix A.

### 3.3 Evaluation metrics

The dice similarity coefficient (*DSC*) is considered as the primary metric in this work. In addition, positive predictive value (*PPV*), sensitivity (*Se*), and the ratio of predicted to target brain volume (*VR*) are also used to evaluate the performance of the brain extraction. These quantities are defined as,

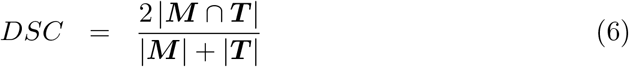

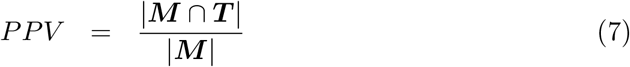

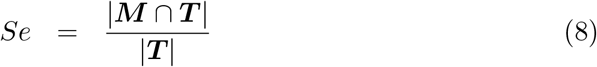

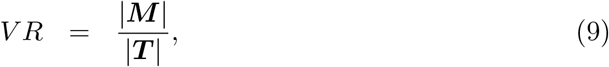

where ***M*** is predicted binary mask, ***T*** is true mask, and | · | denotes the sum of the absolute values of image voxel intensities. The metrics *DSC, PPV*, and *Se* all range from zero to one with the value of one indicating the best performance. Volume ratio (*V R*) can vary from zero to infinity, with one being the optimal value. We note that the entire brain is considered in the calculation of the metrics and no part, like the ventricles, for example, is excluded.

### 3.4 Dataset

We use the Neurofeedback Skull-stripped (NFBS) dataset for the training and evaluating the proposed model. The NFBS dataset consists of 125 anonymized (defaced) T1-weighted 3D MR images of 21 to 45 year old subjects, with a variety of clinical and subclinical psychiatric symptoms. The dataset contains paired images of the head and the brain, where brain extraction is performed using the BEaST method [6] and then corrected manually [22]. This dataset was minimally pre-processed by eliminating voxels with outlier intensities and performing uniform min-max intensity normalization. Each 3D MR image contained *N*_3_ = 170 axial slices, each with *N*_1_ × *N*_2_ = 256 × 192 pixels. Out of the 125 images, 75 (12,750 slices) were used for training and 50 were used for testing the algorithm.

## 4 Results and Discussion

In this section, we present the results of the proposed algorithm as it is applied to a heterogeneous dataset of MR images of the head. Where appropriate, we also discuss the implications of the results.

### 4.1 Brain extraction results

We demonstrate the performance of the proposed algorithm by considering the testing dataset. Figure 5 shows brain extraction results of multiple slices incrementally covering a whole brain of a single patient. In the figure, we start from the top of the head and then progressively move down. From left to right, the first column is the input head image, ***h***, the second column is the ground truth brain image, ***b***, the third column is the predicted pixel-wise mean image, 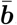, and the fourth column is the pixel-wise standard deviation image, ***b***′. We note that these images have not been post-processed, that is, no thresholding or filtering has been performed.

**Figure 5:**
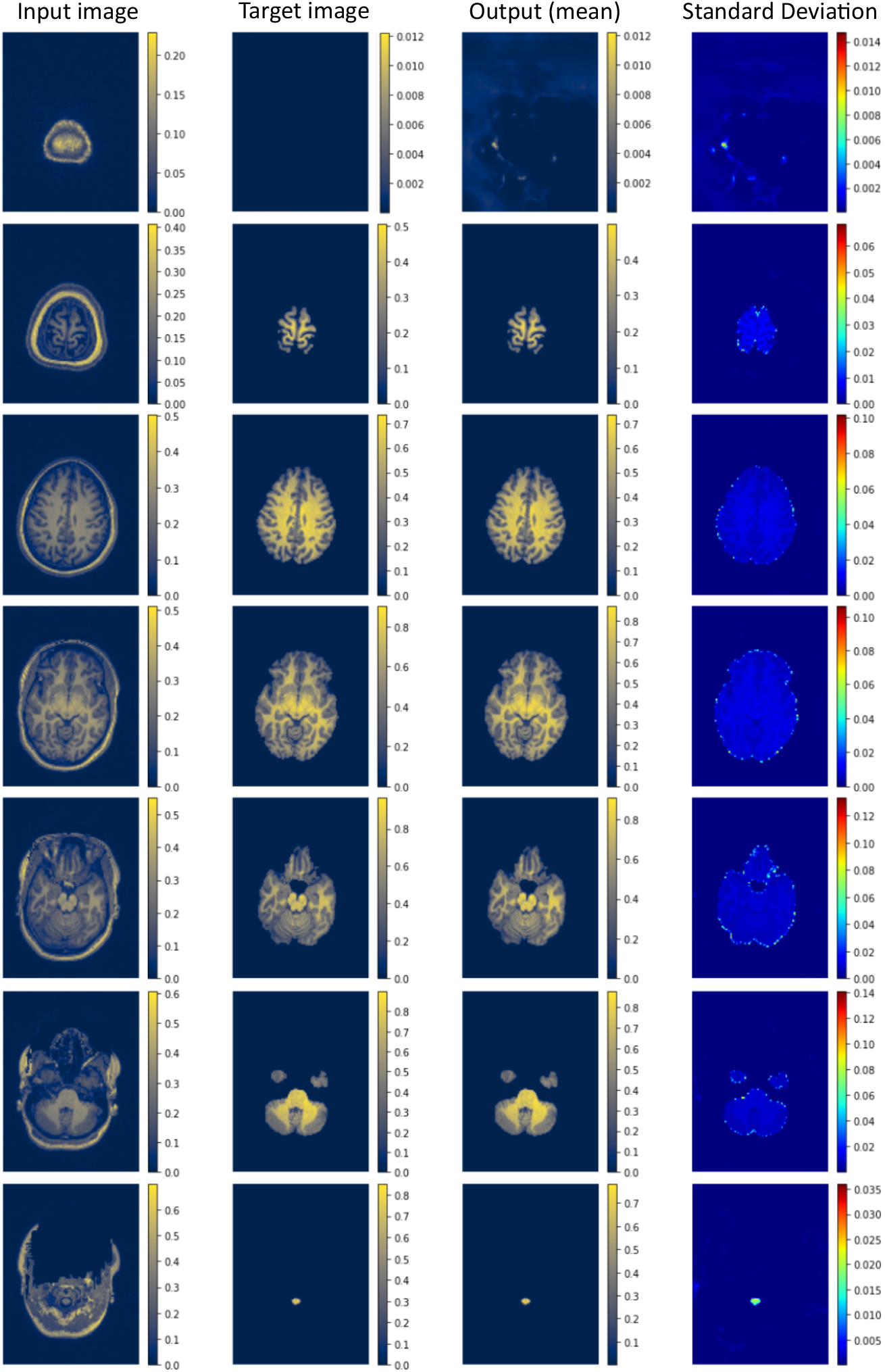
Brain extraction results for a single subject. Left to right: input head image, target brain image, predicted mean brain image, and standard deviation image.

In all slices, we observe that the mean image produced by our algorithm is very close to the target image. Further, the model is robust enough to accurately extract atypical brain parts, such as the brain stem, cerebellum, slices with varied skull thickness, heavily defaced slices, and multi-domain slices where the cerebrum and cerebellum appear in discontinuous locations. In the standard deviation images, we observe that the standard deviation, and therefore uncertainty, is peaked along a thin 2-3 pixel region interface between the brain and the remainder of the image.

A more quantitative assessment of the performance of the model is shown in Figure 6. Here, we evaluate the performance of the binary mask produced by our algorithm (with and without filtering) and the mask produced by the BET tool. We consider two versions of this tool. The first is obtained using the default values of parameters [23]. The second is obtained by performing a grid search to select the “optimal” parameters, which is called BET-optimal (BET-O). In each image, green pixels indicate a true negative prediction (no-brain; correctly predicted), gray pixels indicate a true positive prediction (brain; correctly predicted), orange indicates a false positive prediction (no-brain; incorrectly predicted) and pink indicates a false negative prediction (brain; incorrectly predicted). From left to right, the first column contains the target image and is therefore comprised of only gray and green pixels. The second column is the BET image with default parameters. The third column is the BET image with optimized parameter values, the fourth column is our prediction prior to the application of the filter which eliminates isolated volumes (cGAN), and the final column is our prediction after the application of the filter (cGAN-P).

**Figure 6:**
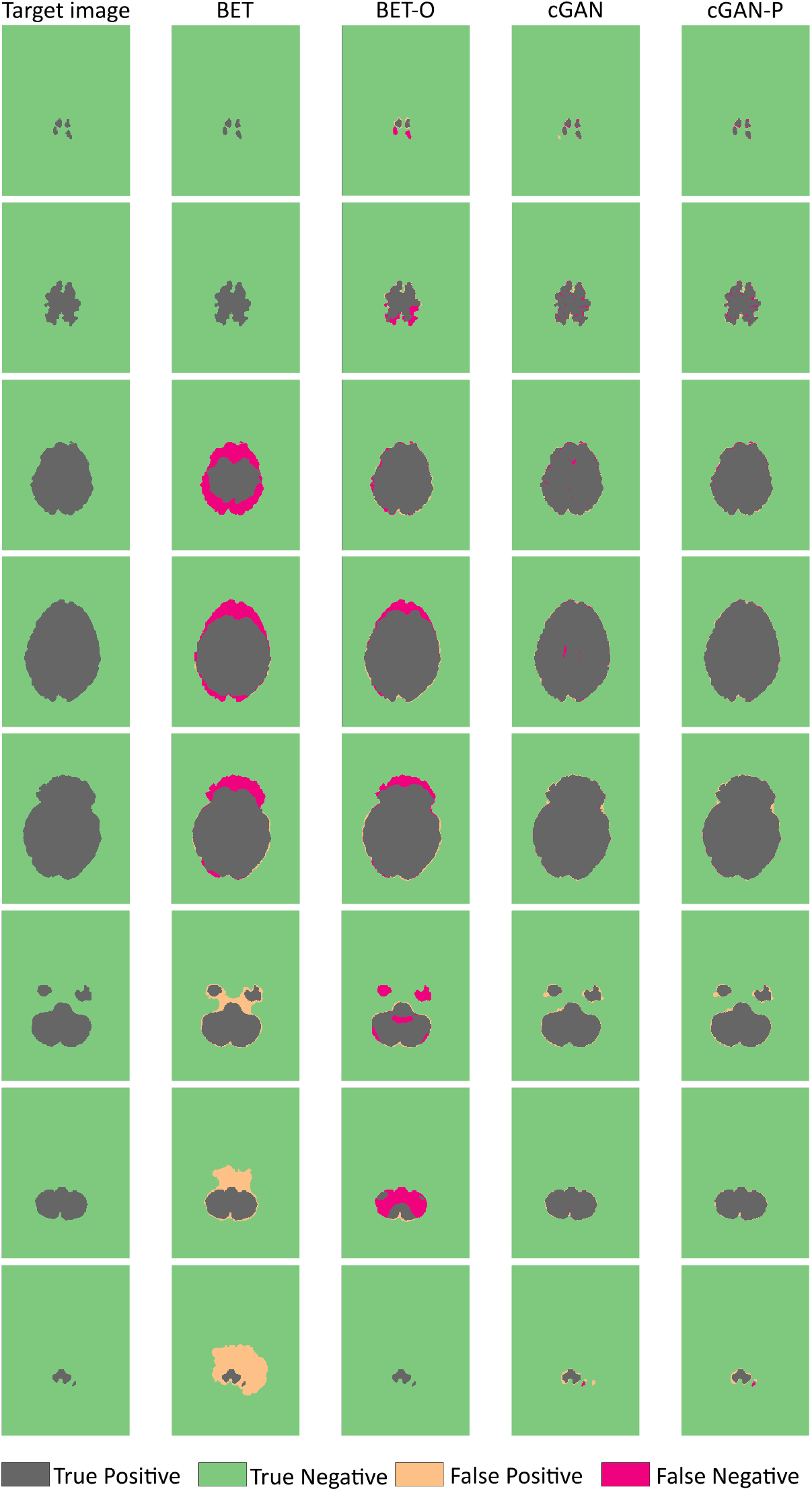
Predicted and ground truth mask image results for multiple slices.

We observe significant differences between the performance of our method and the BET results. The BET method with default parameters (column 2) generates significant regions of false-negative and false-positive predictions. When it is used with optimal parameters (column 3), the false positive regions are reduced, however large false negative regions remain, especially for some of the slices close to the brain stem.

The differences between the results from our method and the BET methods are notable. However, the differences between the two versions of our method (with and without post-processing) are relatively minor. These are most clearly observed in the third and fourth rows, where the application of the volume-deleting filter eliminates the false negative pixels observe in the middle of the brain.

We report quantitative metrics of the performance of the algorithms in Figure 7, where the bars indicate the average value of a metric for the given methods and the whiskers denote the range of the values. These statistics are obtained by applying each of the four methods to the 3D MR images from 50 subjects that were held out for testing. Once again we observe that the conditional GAN approach (with and without post-processing) outperforms the BET-based methods.

**Figure 7:**
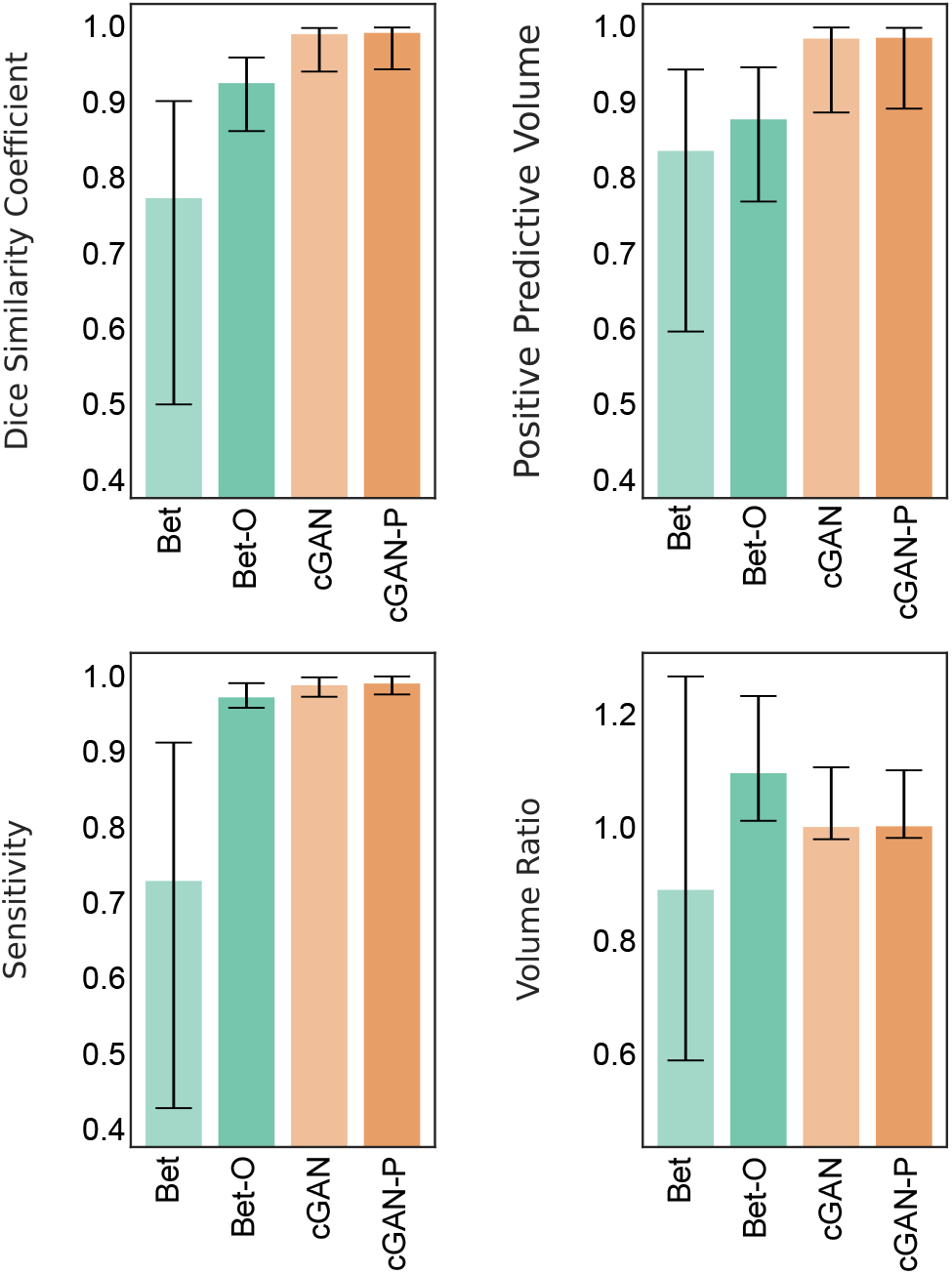
Mean and range of evaluation metrics for BET, BET robust (BET-O), and the proposed model before (cGAN) and after post-processing (cGAN-P).

The improvement obtained from the post-processing step can be observed from Table 1, where we report the mean and the standard deviation (in parenthesis) for the metrics for each method applied to the 50 test images. Note, that we have used a percentage value in this table, and indicated the best-performing methods in bold fonts. We observe that the post-processing procedure nudges the *DSC, PPV* and *Se* values towards perfect scores, and has no discernible effect in altering the prediction of the *V R*.

**Table 1:**
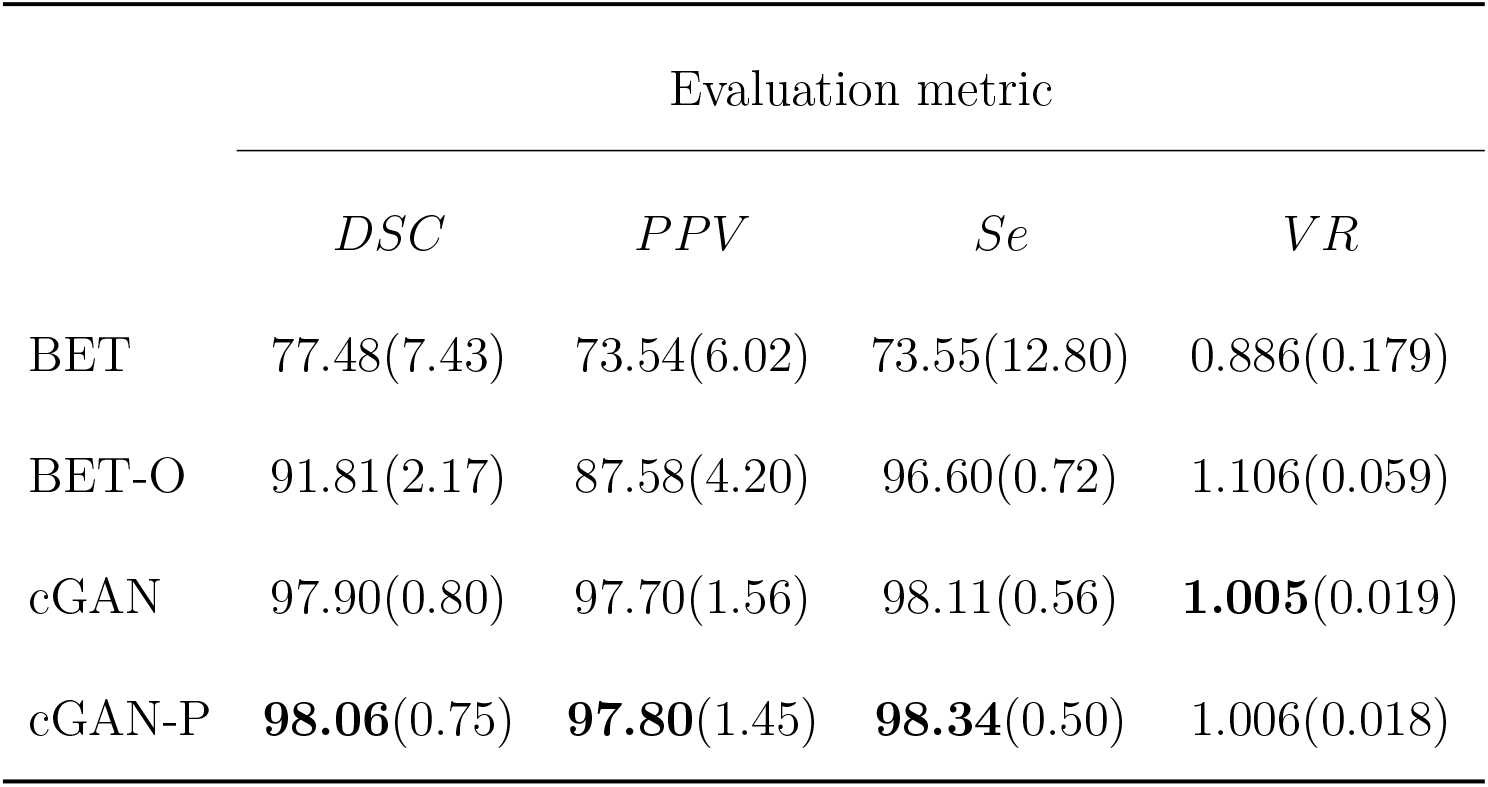
Statistics of evaluation metrics for different methods (in percentage).

|  | Evaluation metric |  |  |  |
| --- | --- | --- | --- | --- |
| | $DSC$ | $PPV$ | $Se$ | $VR$ |
| BET | 77.48(7.43) | 73.54(6.02) | 73.55(12.80) | 0.886(0.179) |
| BET-O | 91.81(2.17) | 87.58(4.20) | 96.60(0.72) | 1.106(0.059) |
| cGAN | 97.90(0.80) | 97.70(1.56) | 98.11(0.56) | **1.005**(0.019) |
| cGAN-P | **98.06**(0.75) | **97.80**(1.45) | **98.34**(0.50) | 1.006(0.018) |

### 4.2 Role of uncertainty quantification

We now discuss how estimates of uncertainty, as given by the pixel-wise standard deviation, can be used to ensure the robustness of a prediction. In Figure 8 we have plotted the head image, the target brain image, the predicted pixel-wise mean, and the predicted pixel-wise standard deviation for one representative head slice. As was seen in Figure 5, once we observe that the standard deviation is typically large at the interface between the brain and the rest of the tissue. However, in Figure 8 it is especially large in a thin region located at the lower-left corner of the image. This region is highlighted by a rectangular box in the standard deviation and the mean images. When comparing the mean and the target images within this region, we recognize that this is precisely where in the mean image our algorithm has incorrectly labeled some pixels as brain (false positives). This example demonstrates that regions of high pixel-wise standard deviation could point the user to regions where the prediction might be incorrect or ambiguous.

**Figure 8:**
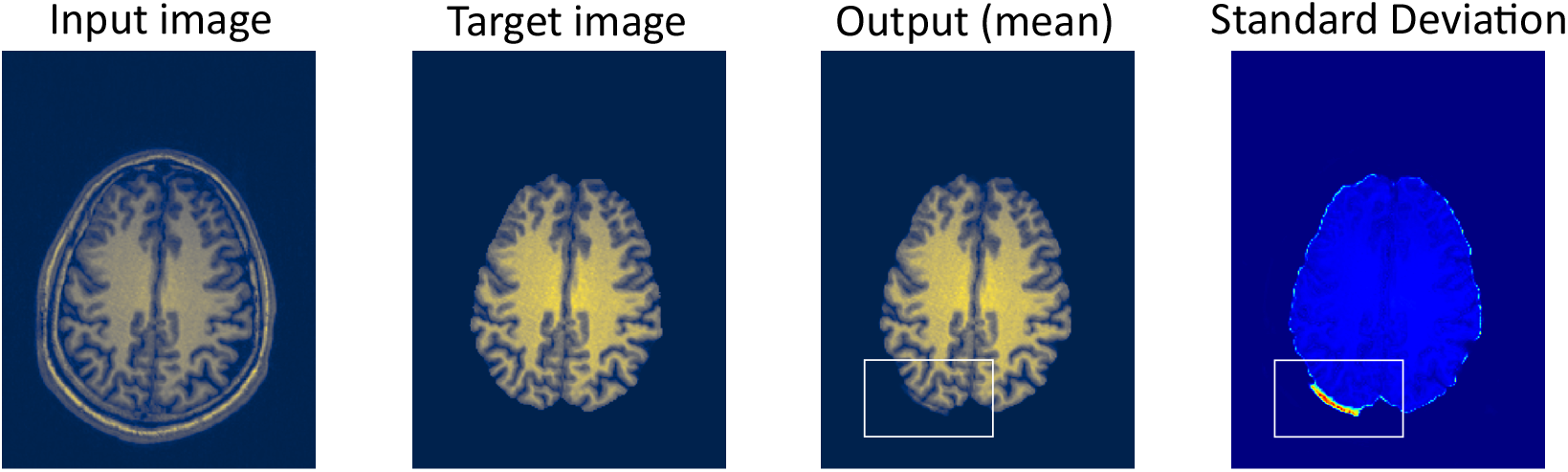
Brain extraction results for a single slice. Left to right: input head image, target brain image, predicted mean brain image, and standard deviation image.

We note that the aggregate value of standard deviation in a 3D image can also be used to assess the performance of the proposed algorithm. This metric can then be utilized by the user to determine how much to trust the reconstructed brain image. This is demonstrated in Figure 9, where for each test subject we have plotted the dice error *ē* = 1 − *DSC* and an aggregate value of the standard deviation. The latter is equal to the sum of the standard deviation of pixels with standard deviation value greater than a threshold (0.04 in this case), divided by the number of axial slices (*N*_3_). In computing this aggregate value, thresholding is performed in order to only include contributions from regions where the uncertainty is large. As can be seen from this figure, there is a strong correlation between the dice error and this measure. We believe that this aggregate measure will be especially useful when the proposed algorithm is used in clinical applications. In those instances, the target brain image and hence the dice error will not be known. However, through our algorithm, the aggregate standard deviation measure will still be calculable. A small value of this measure will provide the user confidence that the resulting brain image is sufficiently accurate.

**Figure 9:**
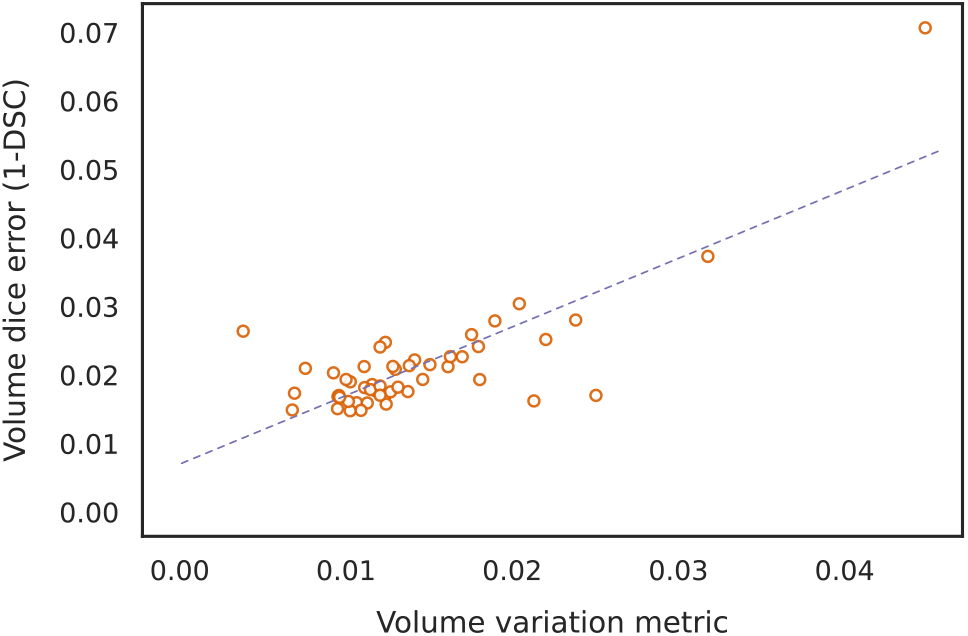
Plot of dice error (1 − *DSC*) and aggregate standard deviation measure for 50 test subjects with a correlation coefficient of 0.805.

## 5 Conclusions

In this manuscript, we have developed, implemented, and tested a novel algorithm for brain extraction that is based on Deep Bayesian inference. It utilizes a conditional GAN formulation, where the generator is in the form of a U-Net and the uncertainty is introduced by the latent variables at multiple scales and for multiple features through conditional instance normalization. The key features of this approach are:

1. Accuracy: The method described in this manuscript yields accuracy metrics that are significantly better than a widely-used brain extraction tool. Further, they compare favorably with the best values reported in the literature by other researchers who have used deep learning and/or U-nets.
2. Uncertainty quantification: to our knowledge, this is the only method in the brain extraction literature that can generate estimates of the uncertainty in the prediction. We also demonstrate how these results can be used to detect regions of likely error within an image, and to assess the overall performance of the algorithm.
3. Speed: the fully trained model can be deployed on a reasonably accessible computer (with Nvidia GeForce RTX 2080 GPU and 16 GB of RAM) and can generate 40 samples in less than a minute (on average 46 seconds over 50 MR images of size 256 × 192 × 170). The BET-O process takes about nine minutes on the same computer.

Future extensions of this work include: (a) Using a 3D U-net architecture to directly process volumes instead of slices; (b) Incorporating this approach in a feedback loop where estimates of uncertainty will be used to improve predictions (with or without human in the loop); (c) Quantifying how the proposed approach to brain extraction can affect the performance of the downstream applications such as brain volume quantification.

## A Neural network blocks

As shown in Figures 3 and 4, the architecture of the generator and critic comprises various network blocks. We describe the key components below.

### Conditional instance normalization (CIN)

The CIN block, depicted in Figure 10(a), is used to inject the latent variable ***z*** into different levels of the generator’s U-Net architecture. It accepts as input ***z*** and an intermediate tensor ***x***^(*i*)^ of size *h* × *w* × *c*. The CIN block first performs a channel-wise normalization (*Norm*) of ***x***^(*i*)^,

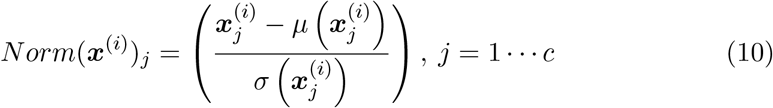

where *μ*(.) and *σ*(.) compute the mean and standard deviation along the spatial directions for a given channel *j*. Next, the latent vector ***z*** of size 1 × 1 × *N*_*Z*_ is passed through two separate 2D convolution layers, ***α***(***z***) and ***β***(***z***), each transforming ***z*** to a tensor of shape 1 × 1 × *c*. The final output of the CIN block is given by the following re-normalization

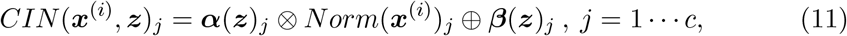

where ⊗ and ⊕ represent element-wise multiplication and summation in the channel direction, respectively. In other words, CIN redefines the channel-wise mean and standard deviation of an intermediate tensor to new values depending (non-linearly) on the latent signal. Note that an advantage of injecting the latent information in this manner is that the dimension *N*_*Z*_ can be chosen independently of the spatial resolution of the MR image.

**Figure 10:**
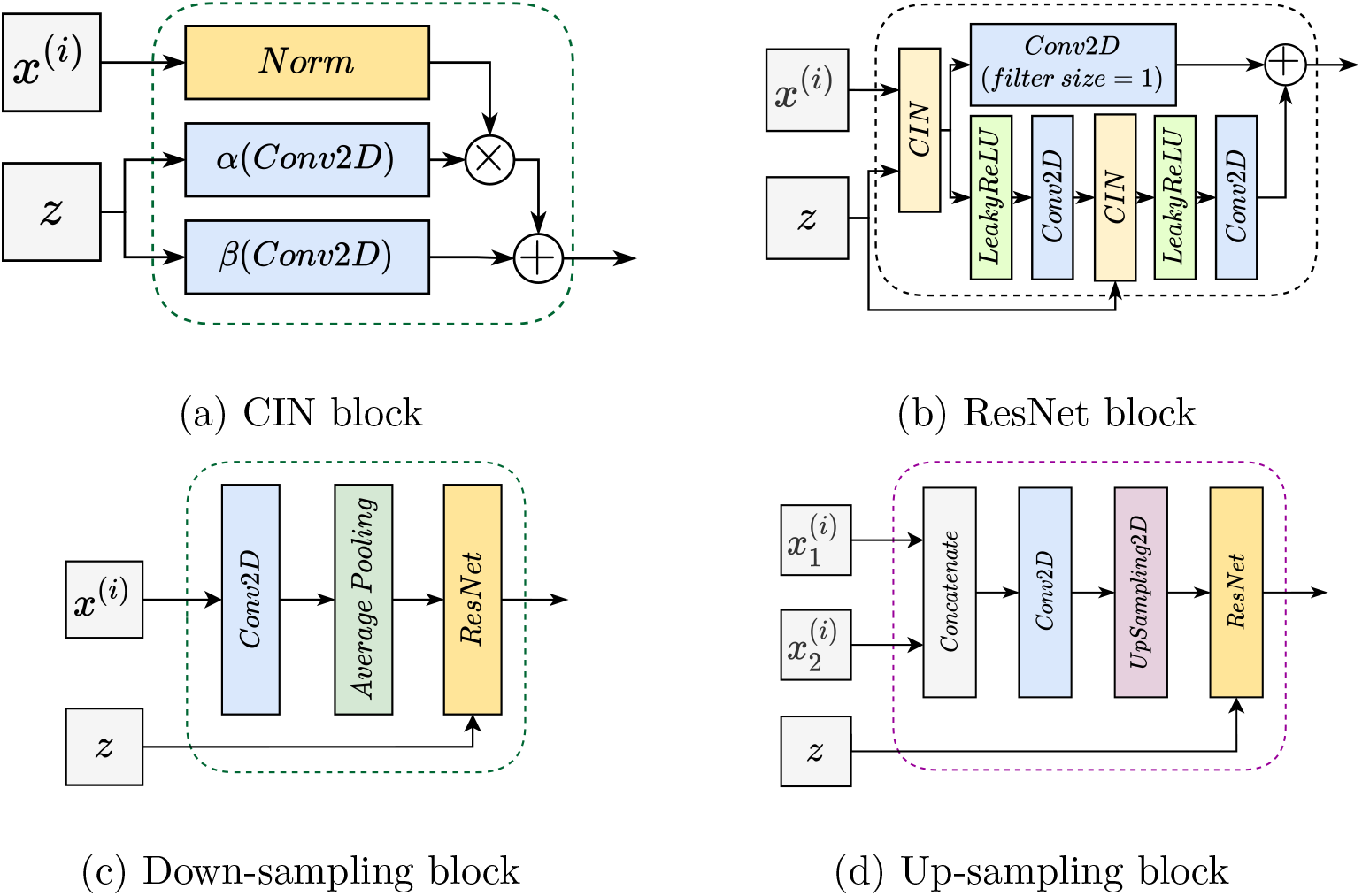
Key components used to build the generator and critic networks.

### ResNet block

Motivated by the network architecture in [15], we implemented a customized ResNet block depicted in Figure 10(b). When appearing in the generator, it takes as input ***z*** and an intermediate tensor ***x***^(*i*)^. The inputs pass through two parallel passways, whose results are summed together to give an output tensor which retains the same shape as ***x***^(*i*)^. Note that when the ResNet block is used in the critic, it takes as input only ***x***^(*i*)^ with CIN replaced by layer normalization.

### Down-sampling block

This block is used to reduce the spatial resolution while increasing the number of channels. As depicted in Figure 10(c), each down-sampling block consists of a convolution layer with output channels twice the number of the channels of the input, a 2D average pooling layer that reduces the spatial dimensions by a factor of two, and a ResNet block.

### Up-sampling block

Each up-sampling block, shown in Figure 10(d), receives the output of the previous block and an output of a down-sampling block of the same spatial size through a skip connection. These tensors are concatenated in the channel dimension. The up-sampling block then performs a convolution that halves the channel size, a 2D up-sampling that increases the spatial dimension by a factor of two, and finally passes the signal through a ResNet block.

